# Phototaxis of the unicellular red alga *Cyanidioschyzon merolae* is mediated by novel actin-driven tentacles

**DOI:** 10.1101/2020.05.09.085928

**Authors:** Sascha Maschmann, Karin Ruban, Johanna Wientapper, Wilhelm J. Walter

## Abstract

Phototaxis – which is the ability to move towards or away from a light source autonomously – is a common mechanism of unicellular algae. It evolved multiple times independently in different plant lineages^1^. As of yet, algal phototaxis has been mainly linked to the presence of cilia, the only known locomotive organelle in unicellular algae. Consequently, phototaxis was believed to be largely absent in red algae (Rhodophyta) that lack cilia in all stages of their life cycle^1^. Remarkably, the unicellular red alga *Cyanidioschyzon merolae* (*C. merolae*) is able to move towards the light. However, it has remained unclear how *C. merolae* can achieve movement, and the presence of a completely new mechanism was suggested^2^.

Here we show that the basis of this movement are novel retractable projections that were termed tentacles due to their distinct morphology. The tentacles could be reproducibly induced within 20 minutes by increasing the salt concentration of the culture medium. Electron microscopy revealed filamentous structures inside the tentacles that we identified to be actin filaments. This is surprising as *C. merolae*’s single actin gene was previously published to not be expressed^3,4^. Based on our findings, we propose a model for *C. merolae*’s actin-driven but myosin-independent motility. To our knowledge, the described tentacles represent a novel motility mechanism.

---

We found that sedimented cells of a liquid culture of the unicellular red alga *Cyanidioschyzon merolae* 10D (*C. merolae*) condensed at a focused light spot within 18 hours (**Fig. 1a,b**). Similar behaviour was already previously described for *C. merolae* and *Cyanidium caldarium*^3^ as well as for *Porphyridium cruentum*^5^. Upon moving the light, the condensed cells immediately started following the spot with a velocity of approximately 2.5 µm s^-1^ (**Fig. 1c, Movie 1**). This movement is considerably slower than swimming phototactic algae like *Chlamydomonas reinhardtii*^6^.

**Figure 1.**
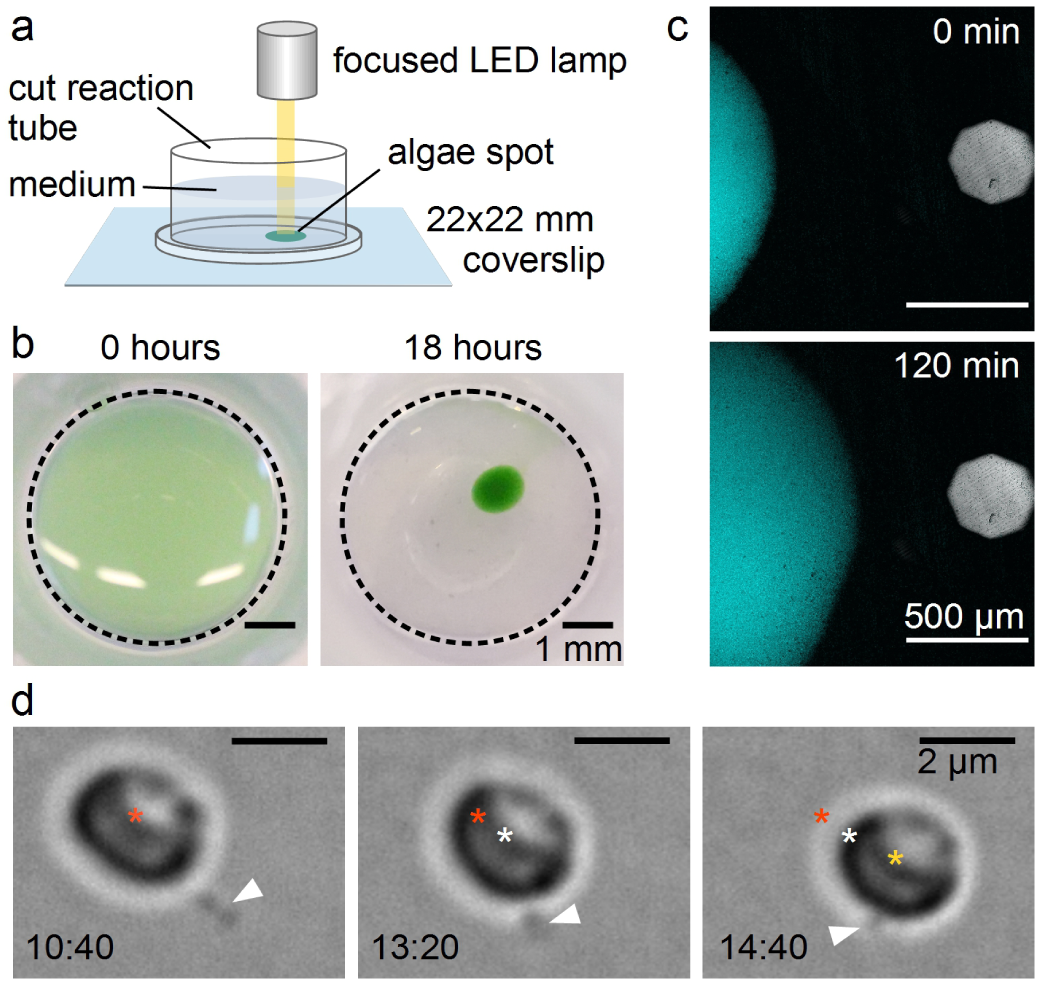
Phototaxis of *Cyanidioschyzon merolae*. **(a)** Set-up for the light microscopical observation of *C. merolae*’s phototaxis. **(b)** Sedimented *C. merolae* culture before and after condensing at a focused light spot. **(c)** Movement of condensed *C. merolae* culture towards a focused light spot. The figure shows a combined autofluorescent signal of the chloroplasts excited at 640 nm (cyan) and a bright field image indicating the position of the focused light spot (grey). **(d)** Single *C. merolae* cell moving by attaching the tip of its tentacle (white arrowhead) to the surface and retracting the projection. Asterisks mark the center of the cell at 0 s (red), 35 s (white) and 70 s (yellow).

Swimming cells are propelled by microtubule-based flagella or cilia^7^ that are lacking in all stages of the Rhodophyta’s life cycle. Some cells, however, can glide over surfaces slowly using pseudopodia facilitated by actin filament dynamics at the cell periphery^8^. *C. merolae*’s genome contains a single actin gene^3,9^. Nevertheless, no actin cDNA clones could be obtained^3^ nor was it detected by fluorescence microscopy with FITC-phalloidin, which specifically binds actin filaments, or by transmission electron microscopy in dividing cells^4^. Consequently, Ohnuma *et al*. proposed a “non-conventional system” to enable *C. merolae*’s movement^2^. Looking at single moving cells, we found that this “system” consists of single or few long projections. Cells attach the tip of the projections to a surface and move forward by retracting it (**Fig. 1d, Movie 2**).

Under optimal growth conditions, *C. merolae* does not form the observed projections, which could explain why they went unnoticed until now. Low light stress induces the formation of projections, yet complicates the observation. Therefore, we tested whether alternative stress factors have a similar effect. We found that addition of 50 mM sodium chloride to the culture medium highly reproducibly induced the growth of mostly one but sometimes up to three projections identical to those formed under light stress (**Fig 2a-c, Movie 3**). The formation of the projections can be divided into two steps. Initially, a bulb appears on the cell’s surface. Subsequently, a thin projection bursts from that bulb, on which the bulb quickly disappears. The projections grow up to a multiple cell length. Using scanning electron microscopy (SEM), we got a more detailed view of the structure of the projections. SEM images show a shank with a diameter of ∼100 nm and an expanded head (**Fig. 2d-f**). We performed transmission electron microscopy (TEM) to get insight into the inner structure of the projections. We found filamentous structures with parallel orientation in the shank and circular orientation in the head (**Fig. 2g-i**). The parallel filament orientation in the shank is highly reminiscent of the parallel orientation of actin filaments in filopodia used by certain animal cells^10^. The circular filament orientation in the head, however, is quite distinct from filopodia and most likely facilitates a sucker-like attachment to the surface by a contraction of the rings. Based on their overall structure and observed function, we decided to term the projections tentacles.

**Figure 2.**
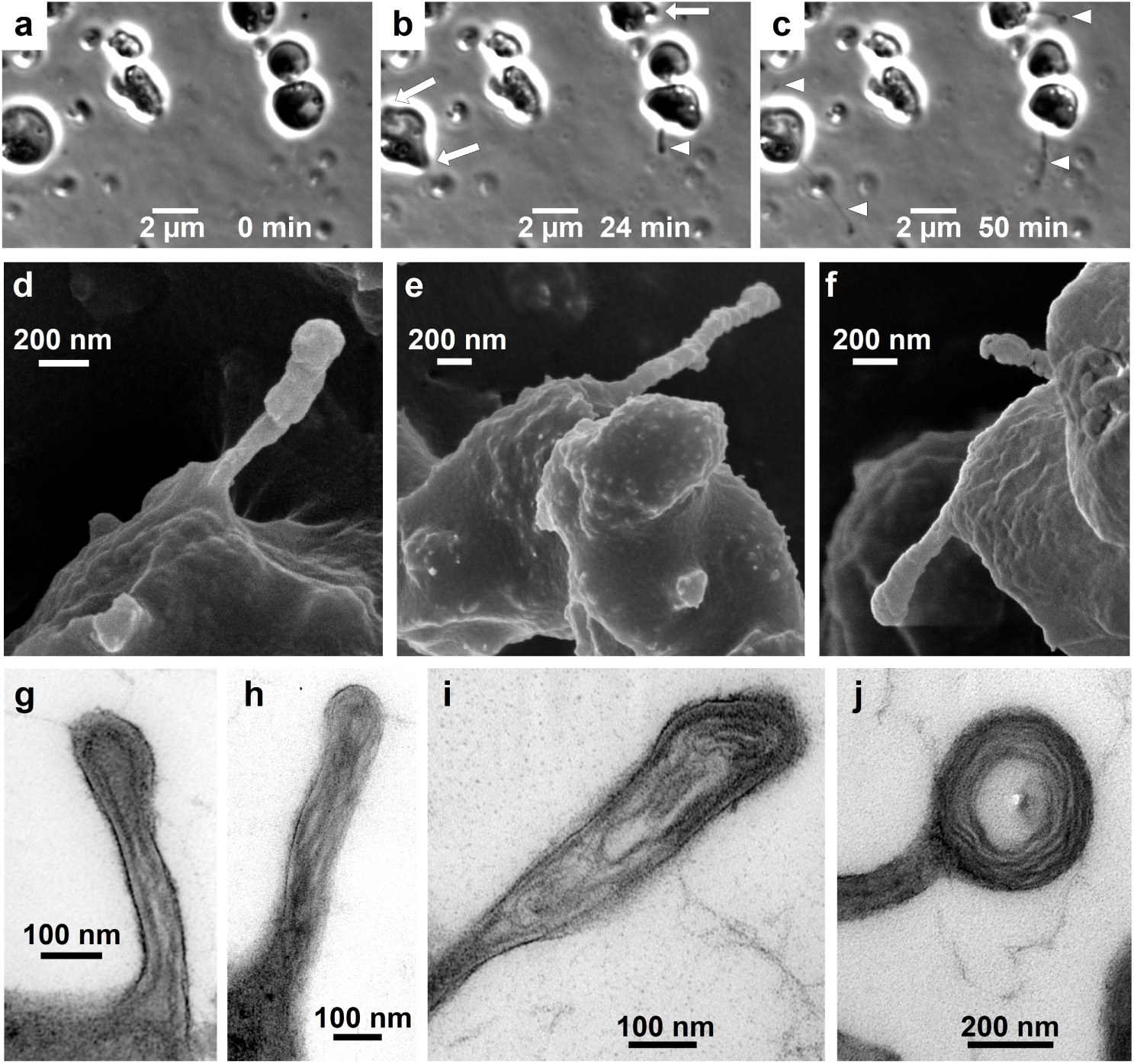
Filopodia-like tentacles in salt-induced *C. merolae* cells. (**a-c**) Time series of *C. merolae* cells growing filopodia-like projections (arrow heads) from bulbs (arrows) upon induction with 50 mM sodium chloride imaged in phase-contrast microscopy. (**d-f**) SEM images show the presence of the extended filopodia-like structures that we termed tentacles. (**g-j**) Negative staining TEM images of thin sections through *C. merolae* tentacles show the presence of long filamentous structures that are oriented in a parallel fashion in the tentacle shaft (g,h) and in concentric circles in the tentacle head (i,j).

Due to the structural resemblance of the observed filaments with F-actin, we decided to re-investigate the expression of *C. merolae*’s actin and actin-related proteins. By alignment of protein sequences using the BLAST algorithm, we identified five actin-related proteins with a sequence identity of 24% to 51% (**Table S1, Fig. S1**). A reverse transcriptase PCR (RT-PCR) with three non-treated and three induced *C. merolae* cultures showed an expression of actin and all actin-related genes independent of the induction (**Fig. 3a**). As this is contradicting previous findings on actin expression in *C. merolae*^3,4^, we performed a Western blot with a monoclonal actin antibody that confirmed our RT-PCR results (**Fig. 3b,c**). As we could demonstrate the expression of actin in general, we stained fixated, salt-induced cells with Alexa Fluor 488 phalloidin. The fluorescence signals co-localized with the tentacles (**Fig. 3d-f**).

**Figure 3.**
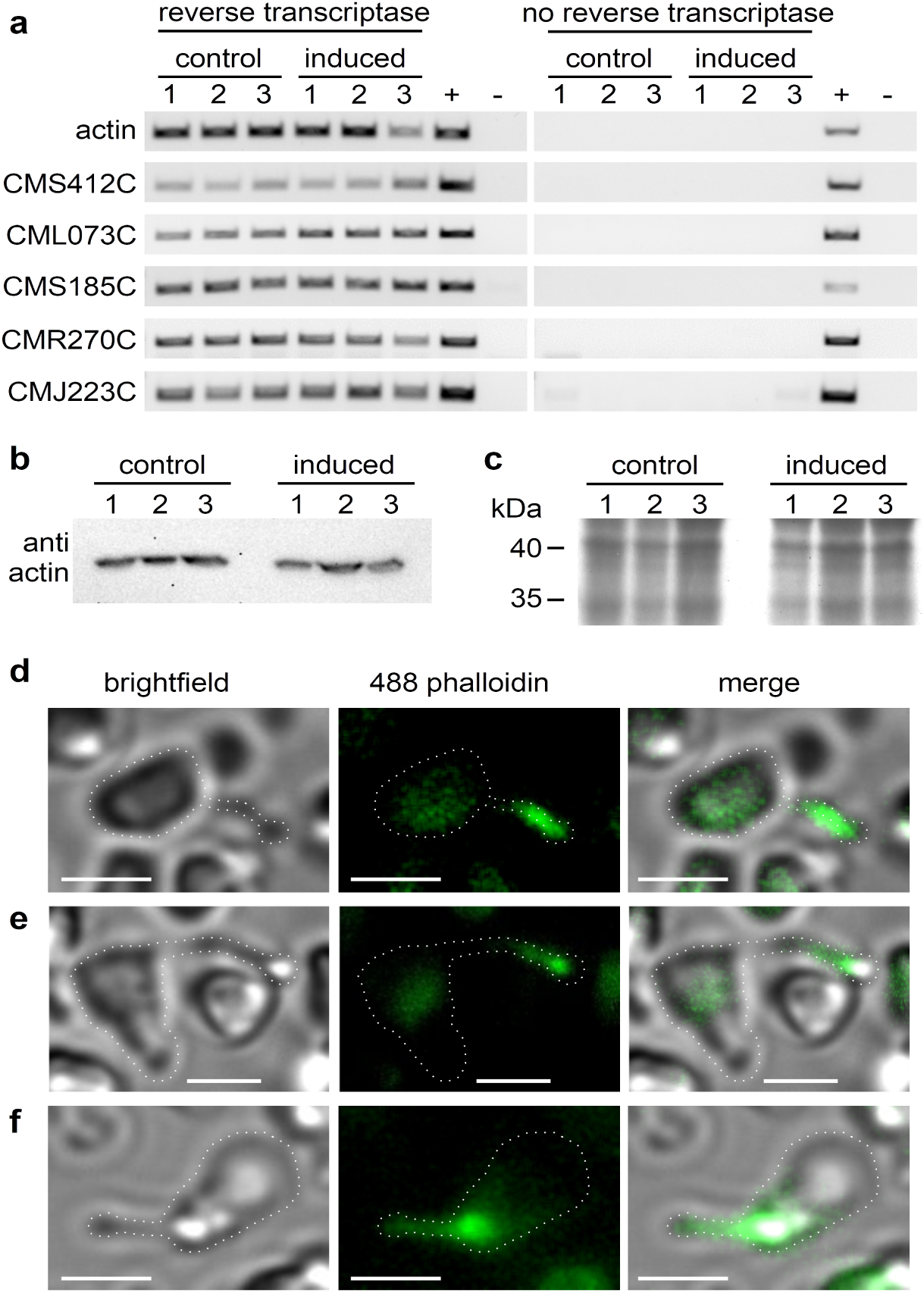
The expression of actin and actin-related proteins in *C. merolae*. **(a)** RT-PCR of of the genes of actin and actin-related proteins before and after tentacle induction. **(b)** Western blot of monoclonal anti-actin antibody clone 10-B3 (Sigma-Aldrich, USA). **(c)** SDS-PAGE of the protein samples used in (b). (**d-f**) Tentacles are positive for staining with Alexa Fluor 488 phalloidin. Brightfield, fluorescence, and merged images of three exemplary cells. Dotted white lines highlight the cell outlines. scale bar = 2 µm

Remarkably, the unchanged levels in actin expression between non-treated and induced cells indicate that tentacle formation in stressed *C. merolae* cells relies on a permanently present actin pool. This pool could explain the rapid formation of tentacles within the first few minutes after induction (**Fig. 2a**).

The tentacles resemble filopodia. Filopodia formation is the result of an orchestrated interplay between actin filaments and multiple well-studied factors including the motor activity of myosins, actin remodelling factors including the ARP2/3 complex, crosslinking proteins such as fascin, α-actinin, or formins, signalling factors like Rho GTPases, and the membrane-deforming activity of members of the I-BAR family^11,12^. However, analogues for most of these proteins, that are essential for the formation of filopodia, are missing in the *C. merolae* genome (**Table S2**). This finding suggests that despite their initial resemblance tentacles are both mechanistically and evolutionarily distinct from filopodia.

Filopodia mostly emerge smoothly from lamellipodia which are sheet-like structures based upon a branched actin filament network^11^. In contrast, we observed that the tentacles burst from a previously formed large bulb (**Fig 2a**). The bulb presumably results from growing actin filaments pushing against a larger plasma membrane area. Unlike in filopodia, the filaments are not guided and assisted by membrane-shaping I-BAR proteins. However, for microtubules, it was shown that the accumulation of crosslinking proteins at the membrane-microtubule interface combined with microtubule polymerization suffices to drive membrane tubulation effectively^12^.

Probably more surprising in the context of an actin-based motile system is the complete absence of myosin motor genes in *C. merolae*’s genome^9^. However, previous studies on filopodia have demonstrated that pulling forces can be exerted independent from myosin. The rearward-directed actin flow, that results from F-actin depolymerization in the filopodia tip, together with inward forces arising from membrane tension is sufficient to generate forces in the low pN range^13,14^. Similarly, the putative attachment of the tentacle head to the surface via contraction of its circular actin structures does not rely on the function of myosins. Recent studies showed a myosin-independent contraction of actin filament rings by the cross-inking protein anillin (https://doi.org/10.1101/2020.01.22.915256). *C. merolae*’s genome does not code for anillin. Nevertheless, other crosslinkers are present and might be able to drive a similar mechanism (**Table S2**).

Besides the mechanical ability to move given by *C. merolae*’s tentacles, phototaxis requires a sensing mechanism that enables an oriented movement response with respect to the direction and intensity of incident light. Other phototactic microalgae like *Chlamydomonas reinhardii* or *Euglena gracilis* can sense the direction of light through a specialized light-sensitive organelle called the eyespot. The eyespot primarily consists of a photoreceptor combined with an arrangement of lipid droplets. The modulation of the light intensity resulting from the respective orientation of the receptor to the lipid droplets allows the cell to adapt its trajectory^15^. Electron microscopy images show that *C. merolae* has multiple arranged lipid droplets potentially serving for sensing light direction (**Fig. S2**). They are less arranged than the eyespot of *Chlamydomonas reinhardtii* which might be due to the significantly smaller cell size and velocity of *C. merolae*. While in green algae, a rhodopsin pigment mediates phototaxis^15^, no rhodopsin-like genes were identified in the *C. merolae* genome. However, several genes coding for blue-light-sensing cryptochromes were found^16^. As *C. merolae* requires a sensing mechanism for phototaxis, it is a fair assumption, that the observed structures play the respective role.

Based on our findings, we propose a novel myosin-independent model for phototaxis in *C. merolae* (**Fig 4**.): (a) Light or salt stress trigger actin polymerization from an existing pool of G-actin. (b) The growing filaments push against the plasma membrane forming the observed bulb. (c) As membrane-shaping I-BAR proteins are missing, a random F-actin bundle prevails to form the growing tentacle. In the tentacle’s shaft parallel actin filaments drive the elongation whereas contraction of circular filaments in the tentacle head allows attachment to the surface. (d) Actin depolymerization and membrane tension enable the retraction of the tentacle. The directionality of *C. merolae*’s movement is governed by a combination of a photoreceptor and an arrangement of lipid droplets.

**Figure 4.**
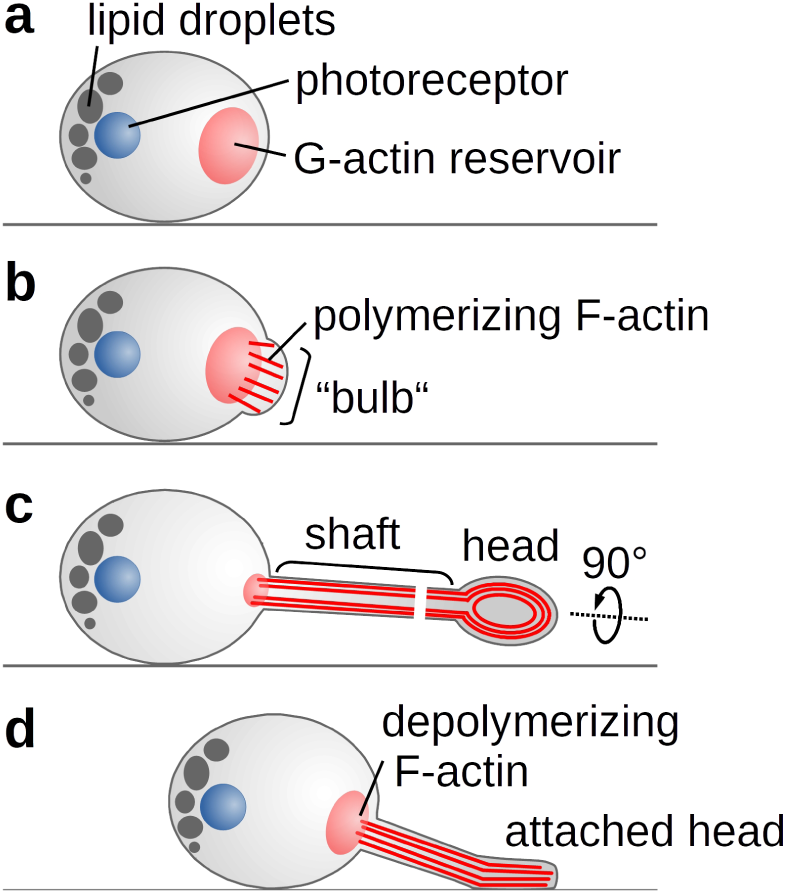
Schematic representation of *C. merolae*’s phototactic mechanism.

To our knowledge, the observed tentacles represent a completely novel organelle for motility. The previously reported two-dimensional phototaxis in other red algae, however, hints towards tentacles being a more general but largely overlooked feature in unicellular Rhodophyta.

## METHODS

Methods and any associated references are available in the supplementary data file.

## ACKNOWLEDGEMENTS

We thank Prof. Dr. Andreas Weber (Heinrich Heine University Düsseldorf, Germany) for providing the *C. merolae* 10D strain as well as Wenke Bahnsen, Elke Woelken, and Renate Walter for technical assistance. We acknowledge support from the Berthold Leibinger Foundation and the Ausgleichststiftung Landwirtschaft und Umwelt to W.J.W.

## AUTHOR CONTRIBUTIONS

W.J.W. designed the experiments, S.M., K.R., J.W., and W.J.W. carried out the experiments; W.J.W. wrote the manuscript. All authors analyzed the data and discussed the results.

## Supplemental information

### Supplemental movies

**Movie 1** | Movement of condensed *C. merolae* culture towards a focused light spot. The movie shows a combined autofluorescent signal of the chloroplasts excited at 640 nm (cyan) and a bright field image indicating the position of the focused light spot (grey).

**Movie 2** | Single *C. merolae* cell moving by attaching the tip of its tentacle to the surface and retracting the projection.

**Movie 3** | *C. merolae* cells growing filopodia-like projections from bulbs upon induction with 50 mM sodium chloride imaged in phase-contrast microscopy.

## Supplemental figures and tables

**Figure S1.**
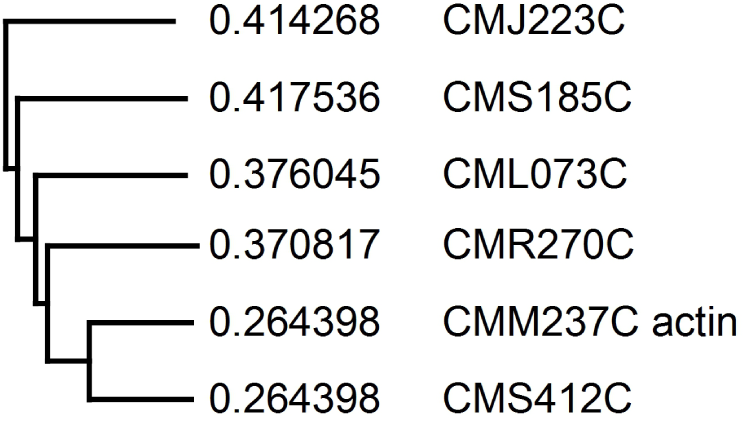
Phylogenetic tree of actin and the actin-related proteins from *C. merolae*.

**Table S1.** Percent identity matrix of actin and the actin-related proteins from *C. merolae*.

|  |  |  |  |  |  |  |
| --- | --- | --- | --- | --- | --- | --- |
| 1: CMM237C actin | 100.00 | 50.93 | 23.78 | 28.57 | 31.01 | 36.17 |
| 2: CMS412C | 50.93 | 100.00 | 25.17 | 27.20 | 29.48 | 31.61 |
| 3: CMJ223C | 23.78 | 25.17 | 100.00 | 21.02 | 27.02 | 27.24 |
| 4: CMS185C | 28.57 | 27.20 | 21.02 | 100.00 | 23.32 | 23.70 |
| 5: CML073C | 31.01 | 29.48 | 27.02 | 23.32 | 100.00 | 27.89 |
| 6: CMR270C | 36.17 | 31.61 | 27.24 | 23.70 | 27.89 | 100.00 |

**Table S2.** Proteins involved in filopodia formation and their analogues in *C. merolae*

|  | **Protein** | ***C. merolae* analogue** |
| --- | --- | --- |
|  | Actin | CMM237C |
|  | Myosin-X <sup>1,2</sup> | none |
| Actin remodelling factors | actin-related proteins <sup>3</sup> | CMS412C, CMJ223C, CMS185C, CML073C, CMR270C |
|  | other ARP2/3 complex proteins (p16, p20, p21, p34, and p40) <sup>3</sup> | none |
|  | cofilin | CMN147C |
|  | profilin | CMP220C |
|  | villin | none |
|  | thymosin | none |
|  | capping protein CP <sup>4</sup> | none |
|  | gelsolin <sup>5</sup> | none |
| F-actin-crosslinking proteins | diaphanous-related formin-2 <sup>6,7</sup> | CMN212C |
|  | formin | CMN049C |
|  | fascin <sup>8,9</sup> | CMD091C |
|  | anillin | none |
| Rho GTPases and other signalling factors | CDC42 <sup>10</sup> | CMJ091C, CMH183C, and multiple putative candidates |
|  | RIF, Rho in filopodia <sup>11</sup> | multiple putative candidates |
|  | WASP/WAVE <sup>12</sup> | none |
|  | ENA/VASP <sup>9,13</sup> | none |
|  | LPR1 , lipid phosphatase-related protein-1 <sup>14</sup> | none |
| I-BAR proteins <sup>15</sup> | insulin receptor substrate p53 | none |
|  | MIM | none |
|  | ABBA | none |
|  | IRTKS | none |

**Figure S2.**
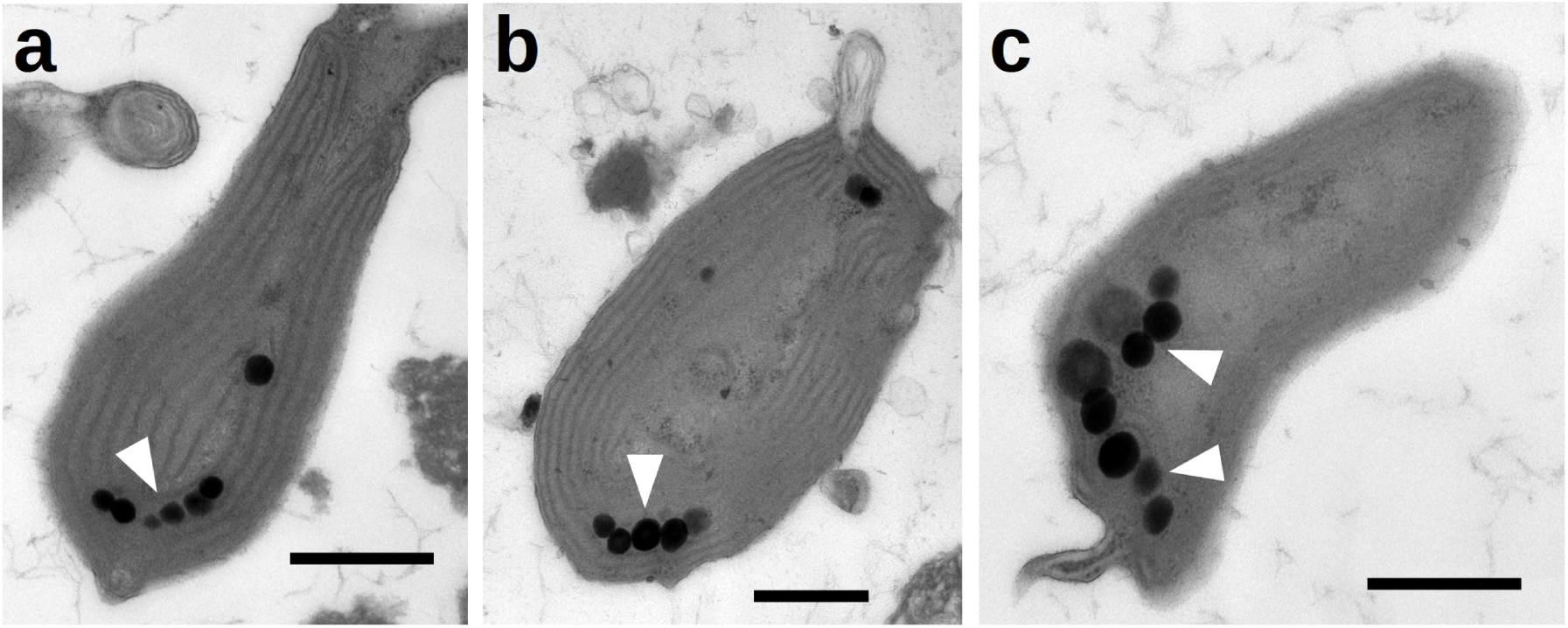
Negative staining TEM of thin sections through three examplary *C. merolae* cells (**a-c**) showing the typical arrangement of lipid droplets. scale bar = 0.5 µm

**Table S3.**
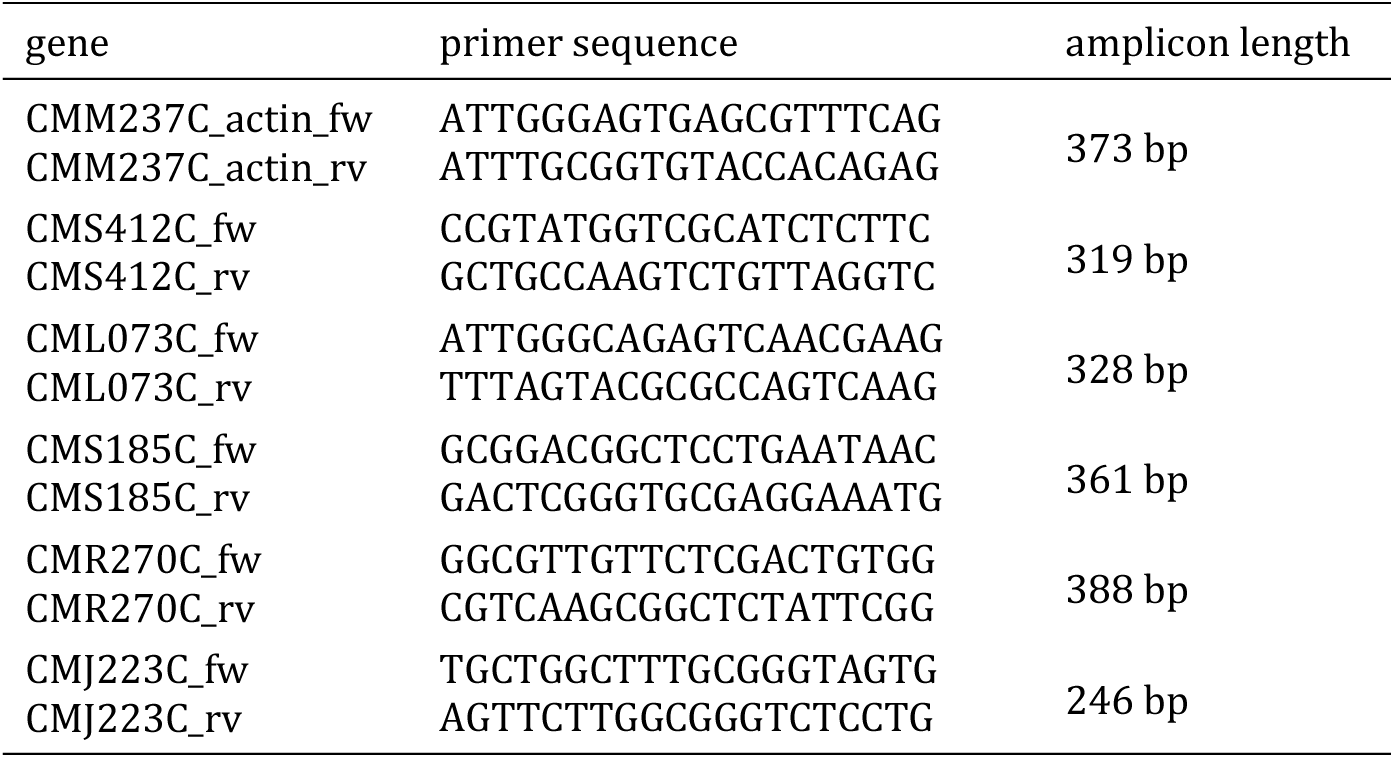
Primers used for the RT-PCR in figure 3

## Methods

### Culture conditions and tentacle induction

The *C. merolae* strain 10D was cultivated in acidic Allen medium (20 mM (NH_4_)_2_SO_4_, 4 mM KH_2_PO_4_, 2 mM MgSO_4_, 1 mM CaCl_2_, 0.3 µM FeCl_3_, 0.7 MnCl_2_, 0.3 µM ZnSO_4_, 70 nM CoCl_2_, 40 nM Na_2_MoO_4_, 10 µM Na_2_EDTA, adjusted to pH 2.3 with H_2_SO_4_) on a heated multiposition magnetic stirrer (RO 15, IKA, Germany) at 300 rpm and 43°C in a 12h/12h day-night cycle with a light intensity of 25 µmol·m^−2^. The addition of 50 mM NaCl induced tentacles. First tentacle formation could be observed after about 15 minutes. After 60 minutes, the majority of cells had formed tentacles (**Fig. 2a**). Control cells were treated with the same volume of water instead of a NaCl solution.

### Actin staining

Samples of a salt-induced *C. merolae* culture were fixated in formaldehyde (4%) for 30 minutes. Actin filaments were stained for 30 minutes by addition of 0.1µm Alexa Fluor 488 (Thermo Fisher Scientific, USA). Cells were gently pelleted and resuspended in Allen medium.

### Light microscopy

Images were acquired by the NIS (Nikon) software packages using an sCMOS camera (Zyla 4.2, Andor, UK) mounted on an inverted fluorescence microscope equipped with an autofocus system (Eclipse Ti, Nikon, Japan). Phase-contrast microscopy images were recorded with the ProgRes CT5 (Jenoptic, Germany). Cells were visualized using bright field, phase contrast, or fluorescence epi-illumination with the respective filter sets.

### Transmission electron microscopy

Samples of an induced *C. merolae* culture were gently pelleted and resuspended in cacodylate buffer (75 mM cacodylate, pH 7.0) supplemented with 2% glutar aldehyde. Samples were fixed for 3.5 hours on ice. Cells were immobilized in 2% agarose in cacodylate buffer before fixing with 1% OsO_4_ over night at 4°C. After washing with cacodylate buffer three times again, samples were dehydrated through a graded series of acetone in cacodylate buffer (30, 50, 70, 90, 100 %) at 4°C with two additional changes in the 100 % at room temperature. Cells were embedded into expoxy resin (ERL-4221D, D.E.R 736, nonenyl succinic anhydride, dimethylaminoethanol) following a protocol by Spurr.^16^

Ultra thin sections (∼80 nm) were prepared on a microtome equipped with a diamond knife (Reichert-Jung Ultracut E) and transferred to copper grids (150 mesh) with a film of polyvinyl butyral (Mowital). The probes were contrasted with solutions of 2% uranyl acetate and 2% lead citrate for 10 minutes each. Images were aquired at 100kV on a transmission electron microscope (LEO 906E, Zeiss, Germany) equipped with a CCD camera (MultiScan 794, Gatan, U.S.A.).

### Scanning electron microscopy

For scanning electron microscopy, samples of a salt-induced *C. merolae* culture were fixated in formaldehyde (1%), dehydrated through an ascending series of ethanol (30, 50, 70, 90, 100 %) and dried at the critical point with Balzers CPD 030 Critical Point Dryer (BALTEC, Germany). After coating samples with gold using a sputter coater (SCD 050, BAL-TEC), scanning electron micrographs were taken with a LEO 1525 (Zeiss, Germany).

### Gene expression analysis

Three induced and three non-induced control samples were frozen in liquid nitrogen and disrupted in a bead mill (Tissue Lyser Qiagen, Retsch GmbH, Germany). Total RNA was extracted from the samples using a purification kit (NucleoSpin, Macherey-Nagel, Germany). For positive controls, samples were not treated with DNaseI. cDNA synthesis was performed using a reverse transcription kit (QuantiTect, QIAGEN, Germany). PCR was conducted using total cDNA as template, gene-specific primers (**Table S3**), and Q5 polymerase (New England Biolabs, U.S.A.) according to manufacturer instructions. Negative controls did not contain template cDNA.

### SDS-PAGE and immunoblotting

Three induced and three non-induced control samples were harvested by centrifugation (2,500 g, 4°C, 60 s). Subsequently the cells were resuspended in extraction buffer (120mM Tris, pH 6.8, 2% SDS, 16% Glycerol, 0.01% Bromphenol blue, 20mM DTT, 0.01% Triton X-100 2×PBS) and heated for ten minutes at 96°C. Cell debris was pelleted (21,000 g, 21°C, 5 min), supernatants were loaded to hand-cast polyacrylamide gels (stacking gel 5% polyacrylamide, 0.1% SDS; resolving gel 10% polyacrylamide, 0.1% SDS). The proteins were stacked at 90 V for five minutes and subsequently separated at 130 V for 45 minutes.

Coomassie staining was conducted according to Dyballa and Metzger^17^.

For immunoblotting, proteins were semi-dry blotted to a PVDF membrane at 3.25 A cm^-2^ for 25 minutes. Subsequently, the membrane was blocked for 1 hour in blocking solution (2% skim milk powder, 1×TBS-T (50mM Tris, 150mM NaCl, 0.05% Tween-20, pH 7.5). The primary anti-actin antibody clone 10-B3 (Sigma-Aldrich, USA) was diluted 1:5,000 in blocking solution and applied to the membrane, slowly shaking at 4°C for 1 hour. After subsequent washing in TBS-T, the secondary anti-mouse IgG peroxidase antibody (Sigma-Aldrich, USA) diluted 1:5,000 in blocking solution was applied for 1 hour at 4°C. Afterwards, the membrane was washed in TBS-T and subsequently, the secondary antibody was detected, by adding chemiluminescence solution (90mM Tris, 227 µg ml^−1^ luminol, 2.06mg ml^−1^ trans-4-hydroxycinnamic acid, 0.014% H_2_O_2_). The chemiluminescence was detected with the ChemiDoc Touch Imaging System (BioRad Laboratories, Germany).

## Notes

### Competing Interest Statement

The authors have declared no competing interest.

